# Cortically-mediated muscle responses to balance perturbations increase with perturbation magnitude in older adults with and without Parkinson’s disease

**DOI:** 10.1101/2024.12.09.627582

**Authors:** Scott E. Boebinger, Aiden M. Payne, Jifei Xiao, Giovanni Martino, Michael Borich, J. Lucas McKay, Lena H. Ting

## Abstract

We lack a clear understanding of how cortical contributions to balance are altered in aging and Parkinson’s disease (PD), which limits development of rehabilitation strategies. Processes like balance control are typically mediated through brainstem circuits, with higher-order circuits becoming engaged as needed. Using reactive balance recovery, we investigated how hierarchical neural mechanisms shape balance-correcting muscle activity across task difficulty in older adults (OAs) with and without PD. We hypothesize that feedback loops involving brainstem and cortical circuits contribute to balance control, and cortical engagement increases with challenge, aging, and PD. We decomposed perturbation-evoked agonist and antagonist muscle activity into hierarchical components based on latency using neuromechanical models consisting of two feedback loops with different delays to reflect different neural conduction and processing times. Agonist muscle activity was decomposed into two components that both increased with balance challenge in both groups. The first component occurred ∼120ms and the second occurred ∼210ms, consistent with the latencies of brainstem and transcortical circuits, respectively. Exploratory comparisons to young adults revealed larger transcortical components in OA and PD groups at lower balance challenge levels, consistent with increased cortical involvement with aging. Antagonist muscle activity included destabilizing and stabilizing components, with the destabilizing component correlating to balance ability in OAs but not in PD. These findings demonstrate that neuromechanical models can identify changes in the hierarchical control of balance without direct brain measurements. Identifying cortical contributions during balance control may complement clinical measures of balance ability to inform balance rehabilitation and assistive devices.

**NEW & NOTEWORTHY:** A neuromechanical model can decompose perturbation evoked muscle activity into components attributed to brainstem and higher-order sensorimotor feedback based on latency. In older adults with and without PD contributions from higher-order neural circuits increase with balance challenge and are related to clinical measures of balance ability. Interpreting the neural substrates of motor output for balance may reveal individual differences in the hierarchical control of balance, that could inform rehabilitation.

## INTRODUCTION

Balance impairments are prevalent in older adults (OAs) with and without Parkinson’s disease (PD) and are associated with less automatic, brainstem-mediated control of balance (1–7), but contributions from higher-order circuits to motor output during balance control remain poorly understood. To maintain standing balance, multisensory information encoding body motion is processed by the central nervous system to generate balance-correcting motor commands to muscles throughout the body (8). Balance control progressively shifts from being primarily brainstem-mediated (9) to engaging higher order centers such as the basal ganglia and cortex as balance health declines, such as in aging and PD (1,10–15), or as balance challenge increases (16,17). Cortical engagement during balance control is typically inferred through measures of cognitive dual task interference (1,10,18–20) or prefrontal cortex oxygen metabolism (13,14,21,22), but these techniques lack the temporal resolution to mechanistically assess the effects on motor output. Reactive balance recovery is a robust paradigm to investigate shifts in sensorimotor control of balance as task difficulty can be adjusted by varying perturbation size. Furthermore, discrete nature of the perturbation and temporal resolution of electromyography (EMG) allows for evoked muscle activity to be attributed to different hierarchical circuits based on latency. Here we investigate shifts in hierarchical motor control by examining muscle activity during reactive balance control using a neuromechanical model to dissociate the contributions of subcortical and cortical components based on feedback delays.

Following external disturbances to either the upper or lower limb, the agonist muscle – i.e. the muscle initially stretched by the perturbation – is activated in a stereotypical sequence of short- and long-latency responses (SLR and LLR, respectively) (23–27). LLRs and are mediated by higher order centers such as the brainstem and cortex, and appear after spinally-mediated SLRs due to a longer neural transmission and processing time (25–29). During balance control, the nervous system maintains the body’s center of mass (CoM) in equilibrium above the base of support (30–32). Following a perturbation to standing, LLRs in the agonist or “prime mover” muscles are patterned based on delayed sensory feedback encoding CoM motion (28,29,33–35). The magnitude and time course of balance correcting agonist muscle activity can be reconstructed based on a small set of feedback parameters that describe the nervous system’s sensitivity to deviations in CoM acceleration, velocity, and displacement (k_a_, k_v_, k_d_) from the desired, upright stance. These scaled CoM kinematic error signals are delayed by a common time delay (λ) to account for neural transmission and processing (28,29,33–35). Prior neuromechanical models used a single sensorimotor feedback loop reflecting brainstem-mediated sensorimotor transforms for the agonist muscle (28,29,33–35).

Recently, we showed balance-correcting muscle activity may reflect feedback components based on multiple sensorimotor transformations from CoM kinematic errors to muscle activity at delays consistent with the involvement of transcortical circuits (15,36). Using a neuromechanical model with parallel sensorimotor feedback loops acting at different delays, we decomposed agonist muscle into components termed LLR1 and LLR2 in young adults (YAs) (36). The LLR1 and LLR2 arise at latencies consistent with brainstem and cortical circuits, respectively (36). In YA, the earlier LLR1 occurred ∼100ms post-perturbation and was engaged at all levels of task difficulty, while the longer-latency LLR2 occurred ∼200ms and became progressively engaged as task difficulty increased (36). We further corroborated that the latency of the LLR2 muscle activity was about twice that of perturbation-evoked cortical activity recorded via electroencephalography (36). Therefore, perturbation-evoked muscle activity may be dissociated into hierarchical components without need for direct brain measurements. However, the dual-loop neuromechanical model has not yet been applied to agonist muscle activity from individuals requiring greater cortical contributions to balance, nor applied to antagonist muscle activity that may also arise from multiple hierarchical neural pathways.

Effective balance recovery requires directionally-specific muscle activation patterns that appropriately generate torque to counteract CoM deviations from an upright stance (37,38). Evoked activity in the antagonist muscle – i.e. the muscle initially shortened by the perturbation – is commonly observed in OAs and individuals with PD, creating destabilizing co-contraction that opposes agonist balance-correcting muscle activity (15,39–45). Antagonist co-contraction increases joint stiffness but reduces net joint torque as the agonist and antagonist muscles resist one another (46). We recently used a neuromechanical model to identify a component of destabilizing antagonist co-contraction activation that was associated with fall history in OAs with and without PD (15). This antagonist co-contraction occurred at latencies ∼ 180 ms but its relative timing with the cortically-mediated LLR2 activity in the agonist muscle has not been investigated.

Here we dissociated hierarchically mediated agonist and antagonist muscle activity in OAs and individuals with PD during control of reactive balance. We hypothesize that parallel sensorimotor feedback loops engaging brainstem and higher-order circuitry contribute to reactive balance control at different latencies and that involvement of higher-order circuits increases with challenge, aging, and PD, increasing recruitment of longer-latency agonist muscle activity and co-contraction of the antagonist muscle (*Figure 1*). We predicted that longer-latency agonist muscle activity would increase with perturbation magnitude and occur at latencies consistent with transcortical mediation in both OAs with and without PD. We further predicted that individuals with PD would have greater longer-latency agonist muscle activity than OAs. Finally, we predicted that the destabilizing antagonist co-contraction and the LLR2 in the agonist muscle are driven by higher- order neural substrates and will therefore occur after the LLR1.

**Figure 1:**
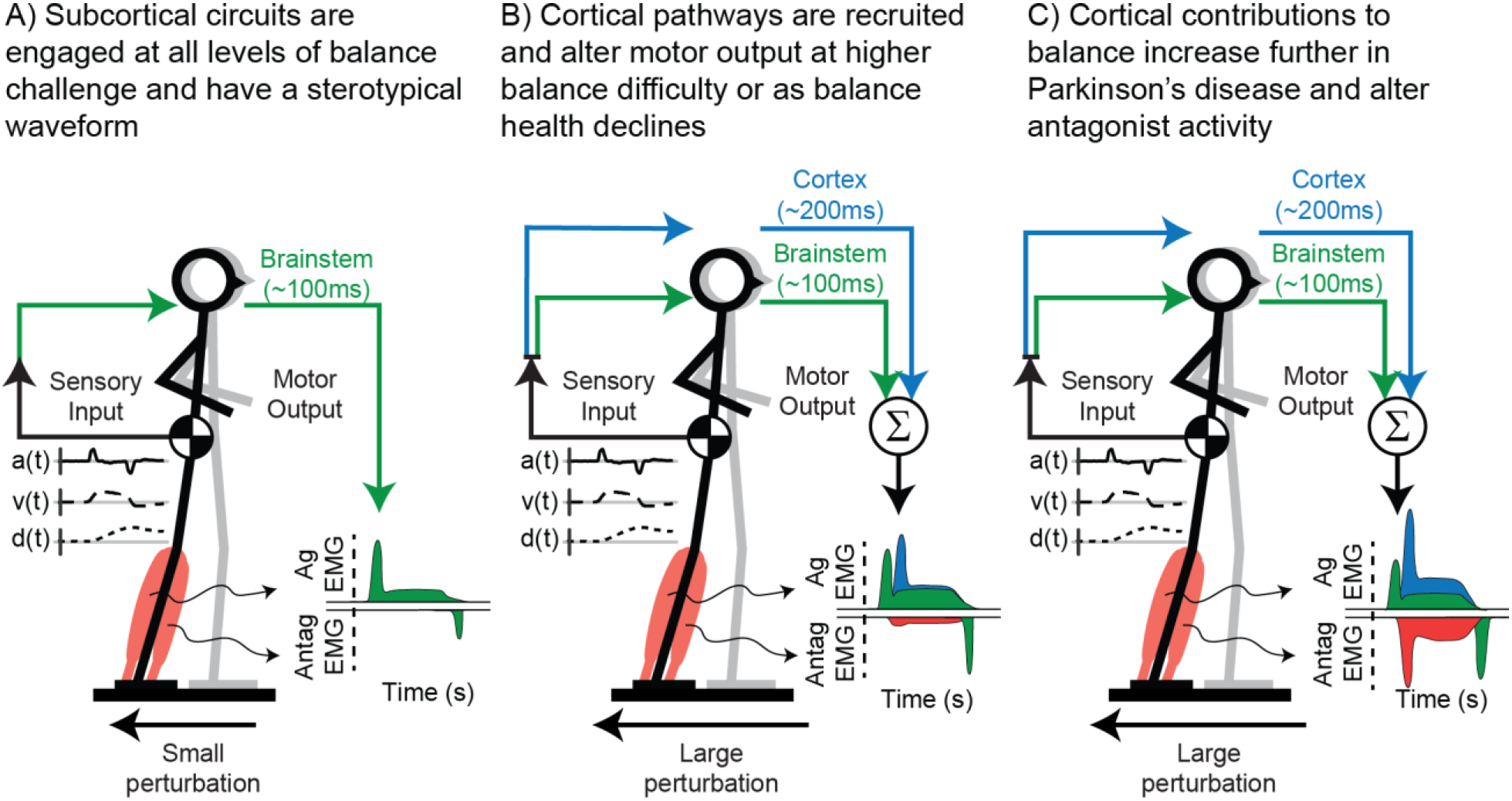
Schematic of hierarchical sensorimotor feedback loops involved in reactive balance control. A) At low levels of balance challenge, balance control is primarily mediated through brainstem sensorimotor circuits (green) which generate a stereotypical EMG waveform. B) At higher levels of balance challenge or as balance health declines, higher order circuits (blue) begin to contribute to balance-correcting muscle activity at longer latencies leading to alterations in the stereotypical waveform. C) In Parkinson’s disease (PD) higher order circuits are further engaged during balance control and leads to abnormal destabilizing antagonist activity (red).

## MATERIALS AND METHODS

### Ethics Statement

All experiments were approved by the Emory University Institutional Review Board and all participants gave informed written consent before participating in this experiment.

### Participants

Nineteen older adults and twenty individuals with PD were recruited for this study. Three participants with PD were excluded from this analysis. Two were excluded due to either a brain tumor or severe peripheral neuropathy of the legs noted in their clinical record. The other opted to leave the experiment prior to balance perturbations. After exclusions, nineteen older adults (6 female, 71 ± 6 years old, 175 ± 2 cm tall, 79 ± 4 kg) and seventeen individuals with Parkinson’s disease (4 female, 69 ± 2 years old, 171 ± 3 cm tall, 84 ± 6 kg) were included in analysis. Participants were excluded prior to participating if they reported having a history of lower extremity joint pain, contractures, major sensory deficits, evidence of orthopedic, muscular, or physical disability, evidence of vestibular, auditory, or proprioceptive impairment, orthostatic hypotension, and/or any neurological insult during recruitment. Participants were recruited from the community surrounding Emory University and Emory Movement Disorders clinic through outreach events, word of mouth, flyers, and databases of prior participants from collaborating groups. Other outcome measures from this group have been reported previously (47–49).

### OFF-medications

Individuals with PD were asked to forgo their dopaminergic medications for PD a minimum of 12 hours before participating in this study. Each participant’s neurologist was consulted and signed a clearance form prior to participants withholding their medications for this study. Clinical and behavioral measures were collected during this OFF-medication session.

### Balance Ability

Participant balance ability was determined via the mini Balance Evaluation Systems Test (miniBEST) which assesses sensory orientation, dynamic gait, anticipatory postural control and reactive postural control (50–52). For items that scored the left and right side separately, only the lower of the two scores was considered which results in a maximum score of 28 (51), where higher scores indicate better balance ability.

### Parkinson’s disease motor symptom severity

For individuals with PD, we examined clinical measures of disease severity to determine if these measures could explain the variance in the hierarchical components identified by our neuromechanical model. The severity of motor impairment in participants with PD was assessed via the motor subscale of the International Parkinson and Movement Disorder Society’s Unified Parkinson’s Disease Rating Scale (MDS UPDRS-III). This test was administered by A.M.P., a certified member of the Movement Disorder Society. Assessments were filmed and sent to a practicing neurologist for scoring.

### Parkinson’s disease duration

The number of years since PD diagnosis was self-reported by participants with PD at the time of the study and verified from their clinical records when possible.

### Balance perturbations

As previously described (47–49), OAs with and without PD stood barefoot on a motorized platform (Factory Automation Systems, Atlanta, GA, USA) and underwent a series of 48 translational support-surface perturbations that were delivered at unpredictable timing, direction, and magnitude. Each participant received an equal number of perturbations in the forward and backward direction at three magnitudes: a small perturbation (5.1 cm, 11.1 cm/s, 0.15 g), which was identical across participants, and two larger magnitudes (medium: 7-7.4cm, 15.2-16.1 cm/s, and 0.21-0.22g, and large: 8.9-9.8cm, 19.1- 21.0cm/s, and 0.26-0.29g). Medium and large perturbations were adjusted based on participant height to ensure perturbations were mechanically similar across participants (47–49) (*Figure 2*). To limit predictability, perturbation magnitude and direction were presented in pseudorandom block order, with each of the eight blocks containing one perturbation of each magnitude and direction. Three different block-randomized orders were used across participants to randomize any effect of trial order (47–49). Trials in which a participant took a step were excluded from our analyses because stepping leaves the position of the base of support temporarily undefined, thereby precluding the calculation of kinematic errors between the CoM and the base of support which are required for our neuromechanical models. Successful non-stepping trials were identified using platform-mounted force plates (AMTI OR6-6). If there were under 3 trials averaged for a certain participant-condition pair it was removed from analysis. To limit fatigue, 5-minute breaks were given every 15 minutes of experimentation. As an additional, exploratory analysis we compared the neuromechanical model outputs from the OA and PD groups to those from a YA group that has been previously published (36). This YA group underwent a similar support surface perturbation paradigm (36,53–55), however there are some key differences to note. Firstly, the YA group underwent larger magnitude support-surface perturbations than the OA and PD groups. Each YA participant received an equal number of perturbations at three magnitudes: a medium perturbation (7.7 cm, 16.0 cm/s, 0.23 g), which was identical across participants, and two larger magnitudes (large: 12.6– 15.0 cm, 26.6–31.5 cm/s, 0.38–0.45 g, and extra-large: 18.4–21.9 cm, 38.7–42.3 cm/s, 0.54–0.64 g), which were adjusted based on participant height (36,54,55). Secondly, the YA group were only perturbed in the backwards direction. To allow for larger perturbations that could sufficiently challenge balance in YAs, the platform was initialized to one end of its range. Due to limitations that the platform can travel, the YA group was only able to undergo perturbations of a single direction while the OA and PD group underwent perturbations in both the backward and forward direction. Due to these differences in experimental paradigm, we only include the YA group as part of an exploratory analysis.

**Figure 2:**
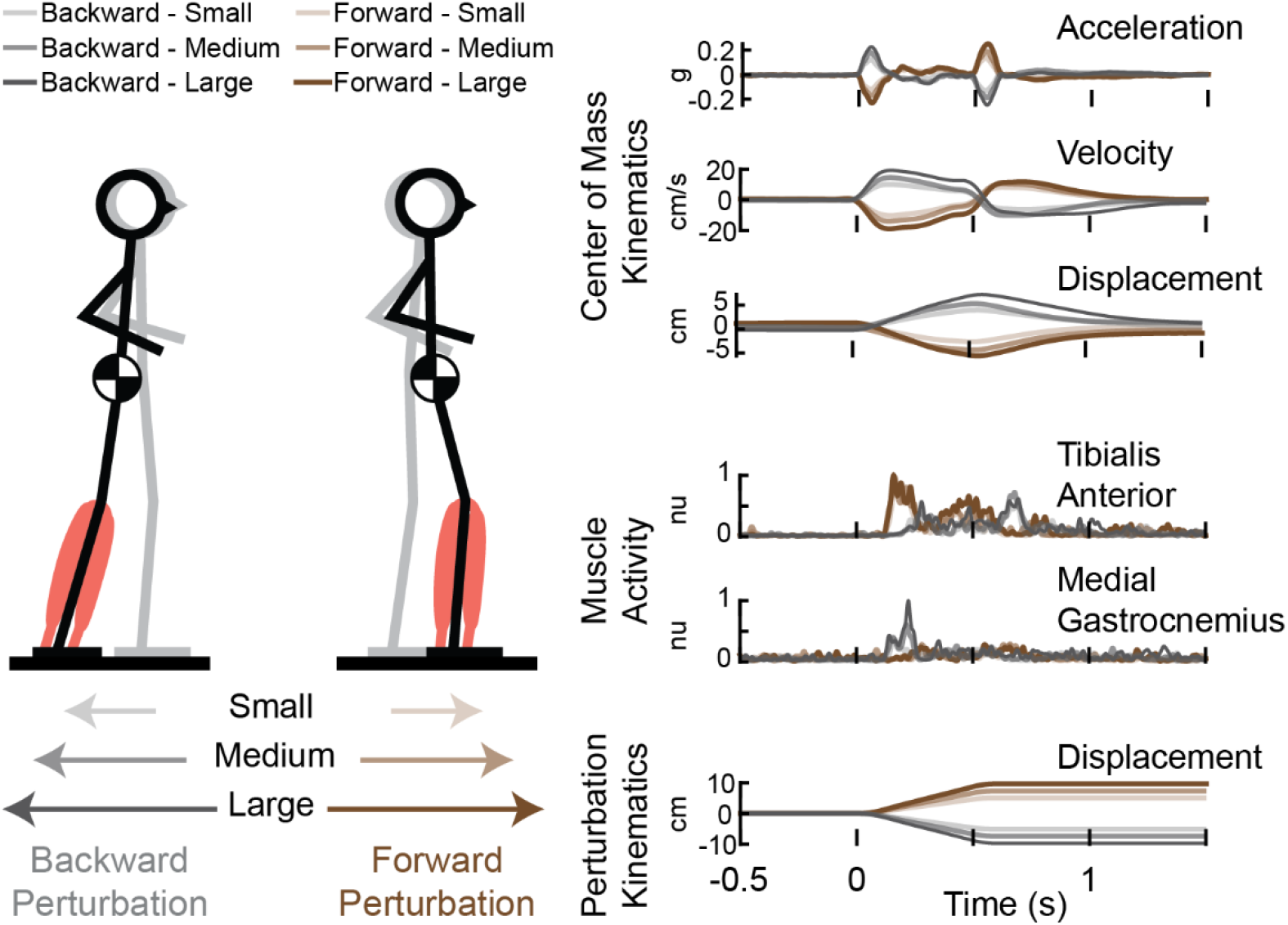
Experimental paradigm. Translational support-surface perturbations were delivered at unpredictable timing, direction, and magnitude. Perturbation kinematics, muscle activity, and center of mass kinematics were recorded throughout the recovery of balance. Brown shaded timeseries represent forward support-surface translations, gray shaded time series represent backward support-surface translations. Darker shading indicates larger perturbation magnitudes.

### Center of Mass (CoM) kinematics

Kinematic marker data were collected at 100 Hz and synchronized using a ten camera Vicon Nexus 3D motion analysis system (Vicon, Centennial, CO). Body segment kinematics were determined from a custom marker set covering head–arms–trunk, thigh, shank, and foot segments. CoM displacement was derived from kinematic data as a weighted sum of segmental masses. CoM velocity was taken as the derivative of CoM displacement after smoothing using a third-order Savtizky-Golay filter with a filter size of 48 samples (35) (*Figure 2*). CoM acceleration was computed from ground reaction forces obtained by the platform-mounted force plates (AMTI OR6-6), divided by participant mass (*Figure 2*).

### Electromyography (EMG)

Surface EMGs (Motion Analysis Systems, Baton Rouge, LA) were collected bilaterally from the tibialis anterior (TA) and medial gastrocnemius muscle (MG) muscles (*Figure 2*). Analysis focused on the TA and MG since this agonist-antagonist muscle pair are activated in forward and backward support surface perturbations. Prior to EMG electrode placement, skin was shaved if necessary and scrubbed with an isopropyl alcohol wipe. EMG electrodes were placed using standard procedures (56). Bipolar silver silver-chloride electrodes were used (Norotrode 20, Myotronics, Inc., Kent, WA, USA). Electromyography signals were sampled at 1000 Hz and anti-alias filtered with an online 500 Hz low-pass filter. Raw EMG signals were then high-pass filtered at 35Hz offline with a sixth-order zero-lag Butterworth filter, mean-subtracted, half-wave rectified, and subsequently low-pass filtered at 40Hz (15,34,55). EMG signals were epoched between -200ms and 1200ms relative to perturbation onset. Single-trial EMG data were normalized to a maximum value of 1 across all trials within each participant for left and right sides independently. EMG data were then averaged across trials within each perturbation magnitude for each participant.

### Neuromechanical models

Perturbation evoked agonist and antagonist muscle activity were reconstructed using a series of delayed feedback models to investigate the relationship between sensory information and muscle activity. All neuromechanical models employed here use kinematic signals of balance error, defined as CoM displacement (d), velocity (v), and acceleration (a) relative to the base of support, as predictors to reconstruct perturbation evoke muscle activity (15,36).

### Agonist neuromechanical model

The agonist neuromechanical model uses two feedback loops with different loop delays to account for the inherently different latencies between brainstem mediated and higher order mediated muscle activity as previously validated in YAs (36) (*Figure 3*). This model is used to decompose perturbation evoked activity in the agonist muscle, defined as the muscle initially stretched by the perturbation, into components attributed to different neural substrates based on latency.

**Figure 3:**
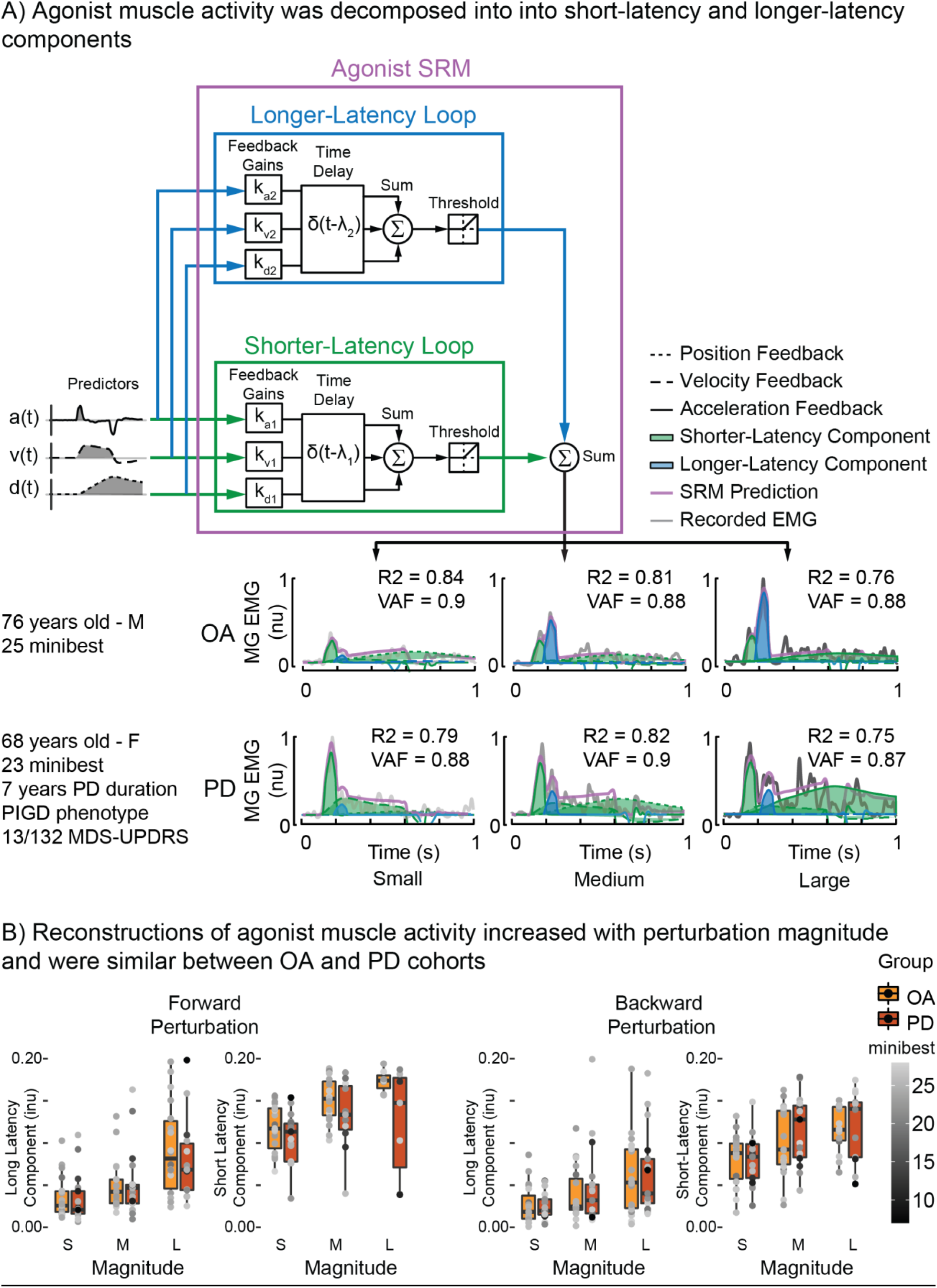
Agonist muscle activity was decomposed into different hierarchical components based on latency_._ A) Agonist muscle activity, defined as the muscle initially lengthened by the perturbation (Medial Gastrocnemius for the backward perturbation shown here), was decomposed into shorter-latency (green) or longer-latency order (blue) components. Agonist muscle activity was reconstructed as the weighted sum of positive CoM kinematics (those that stretch the muscle) that was delayed by two separate time delays reflecting brainstem (λ_1_) or higher order (λ_2_) circuits. B) Group summary of the integrated output of the shorter-latency and longer-latency feedback loops for forward and backward perturbations.

The shorter-latency feedback loop multiplies the CoM predictors that stretch the agonist muscle by their respective feedback gains (k_d1_, k_v1_, k_a1_). These weighted signals are then summed, and delayed by a time delay (λ_1_) to account for ascending and descending neural transmission and processing (*Equation 1*)

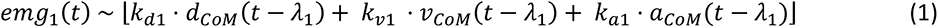

The output from the shorter-latency feedback loop (*emg_1_(t)*) was half wave rectified with a threshold value of 0 to represent the overall non-negative net output to motor pools (*Equation 2*) (15,36).

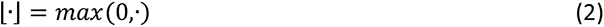

The longer latency feedback loop uses the same predictors that stretch the agonist muscle as the brainstem feedback loop but weights these kinematic signals by independent feedback gains (k_d2_, k_v2_, k_a2_) and delayed by a longer common time delay (λ_2_) to account for the longer latency required for ascending and descending neural transmission and processing in higher-order circuits (*Equation 3*).

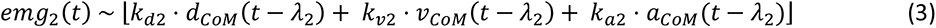

The output from the longer latency feedback loop (*emg_2_(t)*) was also half wave rectified with a threshold value of 0 to represent the overall net output to motor pools (*Equation 2*) (15,36).

The output from both feedback loops were then summed linearly in line with previous modeling approaches of similar behavior in the upper limb (26) (*Equation 4*).

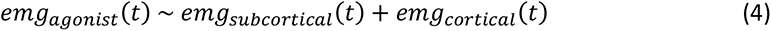

### Antagonist neuromechanical model

The antagonist neuromechanical model also uses two feedback loops with different delays to reconstruct antagonist muscle activity and was developed in a prior study on a separate PD group (15) (*Figure 4*). This model was used to decompose perturbation evoked antagonist muscle activity into stabilizing and destabilizing components.

**Figure 4:**
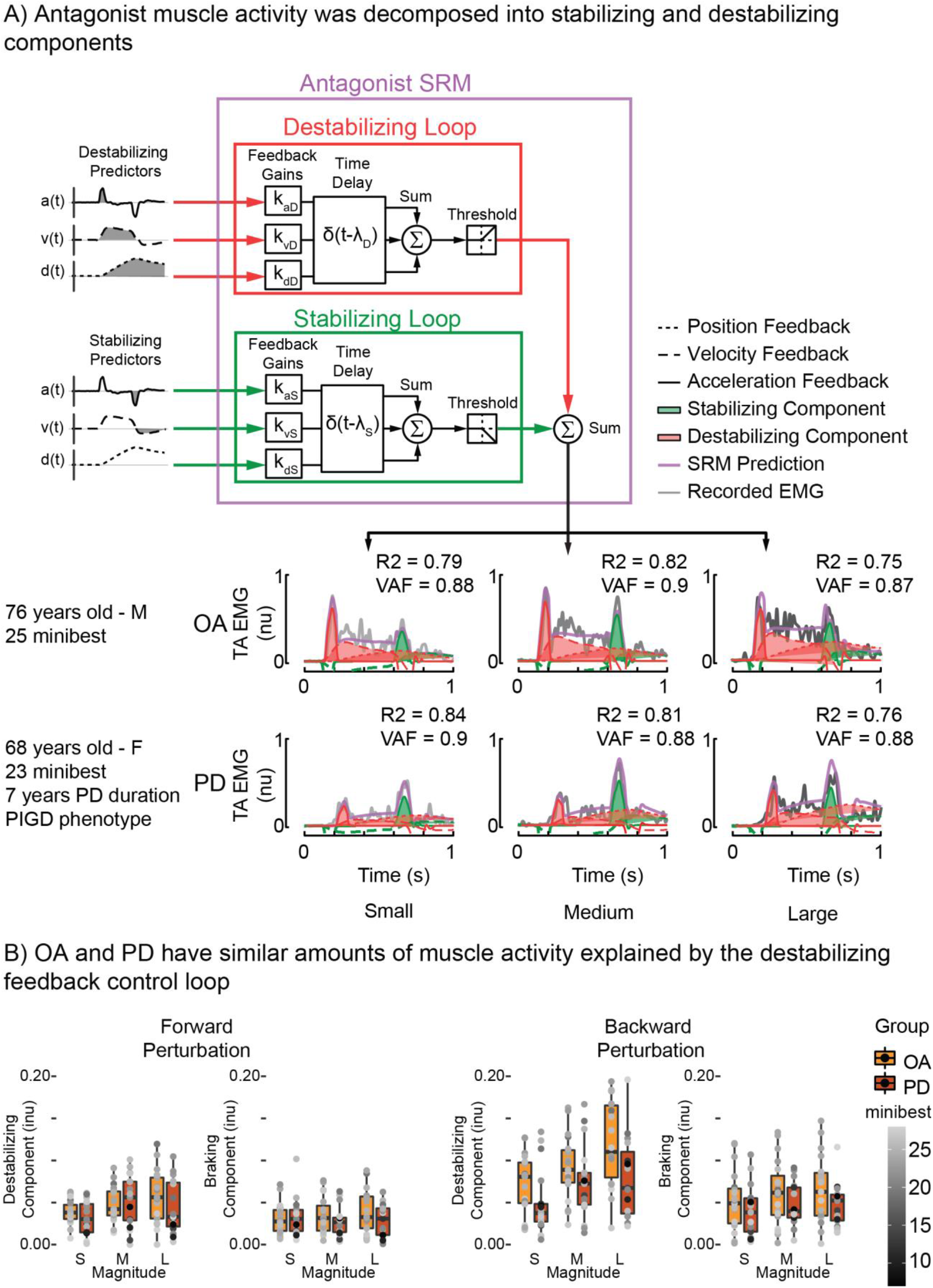
Antagonist muscle activity was decomposed into stabilizing and destabilizing components. A) Antagonist muscle activity, defined as the muscle initially shortened by the perturbation, was decomposed into hierarchical components based off latency using the antagonist neuromechanical model. Destabilizing antagonist muscle activity (red) was reconstructed as the weighted sum of positive CoM kinematics (those that shorten the antagonist muscle) that was delayed to account for neural transmission and processing. Stabilizing antagonist activity (green) was reconstructed as the weighted sum of negative CoM kinematics (those that stretch the antagonist muscle) that was delayed by a separate delay to account for neural transmission and processing. B) Group summary of the integrated output of the stabilizing and destabilizing feedback loops for forward and backward perturbations.

The stabilizing feedback loop multiplies CoM predictors that stretch the antagonist muscle by their respective feedback gains (k_dS_, k_vS_, k_aS_). These weighted signals are then summed, and delayed by a common time delay (λ_s_) to accounts for ascending and descending neural transmission and processing (*Equation 5*)

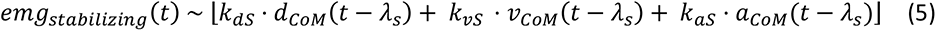

The output from the stabilizing feedback loop (*emg_stabilzing_(t)*) is half wave rectified with a threshold value of 0 (*Equation 2*).

The destabilizing feedback loop uses CoM predictors that shorten the antagonist muscle (-a(t), - v(t), and -d(t)) and multiplies these kinematic signals by separate feedback gains (k_dD_, k_vD_, k_aD_). These weighted signals are them summed and delayed by a common time delay (λ_D_) to account for ascending and descending neural transmission and processing (*Equation 6*).

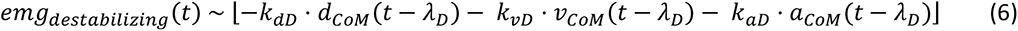

The output from both the destabilizing (*emg_destabilizing_(t)*) feedback loops was also half wave rectified with a threshold value of 0 to represent the overall net output to motor pools (*Equation 2*) (15).

The output from both feedback loops were then summed linearly similar to the reconstructed agonist activity (*Equation 7*).

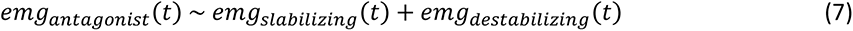

### Model parameter identification

Parameters for both the agonist and antagonist neuromechanical models were selected by minimizing the error between recorded EMG data and the model reconstruction. The reconstruction error was quantified as the sum of the sum squared error of the whole time series as well as the maximum error observed at any sample (*Equation 8*).

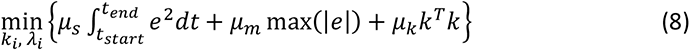

The first term penalizes squared error (e^2^) between the recorded data and the model prediction weighted by μ_s_. The second term penalizes the maximum error observed between the recorded data and the model reconstruction weighted by μ_m_. The final term penalizes the magnitude of the gain parameters (k_i_) with weight μ_k_ in order to minimize feedback parameters that do not contribute to accurate reconstructions. The ratio of weights μ_s_:μ_m_:μ_k_ was 1:1:1e-6. All optimizations were performed in Matlab 2022b (Mathworks, Natick, MA, USA) using the interior-point algorithm in *fmincon.m*, as described previously in literature (15,36).

Separate optimizations were performed to identify model-specific feedback parameters for the individual feedback loops (*Equations 1, 3, 5,* and *6*). The output from these optimizations are then used as an initial guess when optimizing for double-looped models (*Equations 4* and *7*) similar to what was done in prior implementations of this model (15,36). This allows for modifications to the single-loop optimization’s output, as model parameters may have been overestimated in order to fit muscle activity that is better explained by the other feedback loop. Lower and upper bounds for the gain parameters were ±10% of the initial guess values, lower and upper bounds for the delay parameters were ±10ms of the initial guess values. In all cases, additional parameters supplied to *fmincon.m* were as follows: *TolX*, 1e-9; *MaxFunEvals*, 1e5; *TolFun*, 1e-7, with remaining parameters set to default (15,36).

### Goodness of fit

The goodness of fit for all neuromechanical model reconstructions were assessed using a coefficient of determination (R^2^) and variability accounted for (VAF). R^2^ was calculated using *regress.m*, a built-in function in Matlab 2022b (Mathworks, Natick, MA, USA). VAF was defined as 100*the square of Pearson’s uncentered correlation coefficient, as performed in previous studies (15,36,57).

### Model selection

Model selection was performed using Akaike’s Information Criterion (AIC) (*Equation 9*).

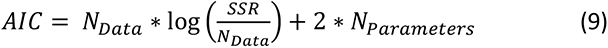

N_Data_ is the number of data points reconstructed by the neuromechanical models (1400 samples), SSR is the sum squared error of the neuromechanical model reconstruction, and N_Parameters_ is the number of parameters included in the neuromechanical model (8 parameters). AIC values were calculated for both the agonist and antagonist neuromechanical models described above. An AIC comparison was performed to determine whether the addition of a longer-latency loop improves the model fit to the data in the agonist neuromechanical model. Differences in AIC between models ≥ 2 were considered meaningful.

### Statistical characterization of neuromechanical model outputs

The integrated area under the curve of each neuromechanical model component (*Equations 1, 3, 5,* and *6*) was calculated via numerical integration using *trapz.m*, a built-in function in Matlab 2022b (Mathworks, Natick, MA, USA). Statistical tests were performed in RStudio version 1.4.1717 (R Core Team, Vienna, Austria). Comparisons of integrated components between perturbation magnitude and group were performed using a linear mixed effects model with an interaction between the fixed factors of perturbation magnitude and group with participant as a random factor. Post hoc comparisons were performed via comparisons of the estimated marginal means between perturbation magnitudes or group. Correlations between integrated components and clinical measures (miniBEST and MDS UPDRS-III scores) were performed using a linear mixed effects models with interaction between the fixed factors of clinical score and perturbation magnitude with participant as a random factor. Separate linear mixed models were applied for the OA and PD groups. Tests are considered statistically significant at p ≤ 0.05.

## RESULTS

### Agonist sensorimotor response to perturbations

Perturbation evoked agonist muscle activity exhibited an LLR1 followed by an LLR2 occurring around 100 and 200 ms latency, respectively, in both OA and PD groups. Both the LLR1 and LLR2 could be explained by sensorimotor transformations of CoM kinematics via our double looped neuromechanical model (*Figure 3A*). The shorter-latency feedback loop (*Figure 3A*, green trace) explaining the initial LLR1 of agonist muscle activity occurred at 119ms ± 1.6ms for OAs and 116ms ± 1.3ms for individuals with PD, with no differences between groups in forward (F(1, 34) = 1.87, p = 0.18) or backward perturbations (F(1, 34) = 0.02, p = 0.89). The longer-latency feedback loop (*Figure 3A*, blue trace) reconstructing the LLR2 of agonist muscle activity occurred at 200ms ± 3.2ms in OAs and 215ms ± 3.9ms in individuals with PD, with a significant differences between groups in backward perturbations (Forward: F(1, 33.9) = 0.46, p = 0.50; Backward: F(1, 34) = 4.17, p = 0.049). Goodness of fit of agonist neuromechanical model reconstructions increased by ∼10% when using a double versus single feedback loop model in both groups (*Table 1)*. This led the double loop model to have higher reconstruction accuracies compared to the single loop model in both perturbation directions (Forward: R^2^: t(34.5) = 11.2, p < 0.0001; VAF: t(34.7) = 11.0, p < 0.0001; Backward: (R^2^: t(35) = 6.6, p < 0.0001; VAF: t(35) = 6.3, p < 0.0001).

**Table 1:** Reconstruction accuracies for single loop and double loop agonist neuromechanical model.

| Model Type | Older Adults | Individuals with Parkinson's disease |
| --- | --- | --- |
| Single Loop Model | $R^2 = 0.64 \pm 0.01$<br>$\text{VAF} = 0.74 \pm 0.01$ | $R^2 = 0.61 \pm 0.02$<br>$\text{VAF} = 0.72 \pm 0.13$ |
| Double Loop Model | $R^2 = 0.74 \pm 0.01$<br>$\text{VAF} = 0.83 \pm 0.01$ | $R^2 = 0.71 \pm 0.01$<br>$\text{VAF} = 0.81 \pm 0.01$ |

Values are mean ± SEM.

Contributions of both the LLR1 and LLR2 increased with perturbation magnitude in both OA and PD groups, with no difference between groups (*Figure 3B*). In both perturbation directions, there was an increase with perturbation magnitude in the LLR1 (Forward: F(2, 65.8) = 36.24, p < 0.001; Backward: F(2, 68) = 25.98, p < 0.001) and LLR2 (Forward: F(2, 64.7) = 37.6, p < 0.001; Backward: F(2, 68) = 16.97, p < 0.001). There was no difference between OA and PD groups in either the LLR1 (Forward: F(1, 34.6) = 0.37, p = 0.55; Backward: F(1, 34) = 0.31, p = 0.58) or LLR2 (Forward: F(1, 34.6) = 0.08, p = 0.78; Backward: F(1, 34) = 0.12, p = 0.73) (*Figure 3B*). There were no interactions between perturbation magnitude and group for either the LLR1 (Forward: F(2, 65.8) = 1.36, p = 0.26; Backward: F(2, 68) = 0.22, p = 0.81) or LLR2 (Forward: F(2, 64.7) = 0.06, p = 0.94; Backward: F(2, 68) = 0.34, p = 0.71).

### Antagonist sensorimotor response to perturbations

Perturbation-evoked antagonist muscle activity could also be explained by sensorimotor transformations of CoM kinematics via a double looped neuromechanical model (*Figure 4A*). Antagonist muscle activity was exhibited destabilizing co-contraction at the beginning of the ramp and hold perturbation that could be explained by sensorimotor transformations of CoM kinematics in the destabilizing feedback loop (Figure 4A, red trace). This co-contraction was then followed by a stabilizing response to platform deceleration that could also be explained by the sensorimotor transformation of CoM kinematics in the stabilizing feedback loop (Figure 4A, green trace).

Contributions of the destabilizing feedback loop to antagonist muscles (*Figure 4A* red trace) increased with perturbation magnitude in both OA and PD groups (Forward: F(2, 63.8) = 10.15, p < 0.001; Backward: F(2, 68) = 44.12, p < 0.001) (*Figure 4B*). There were no differences between the OA and PD groups in either perturbation direction (Forward: F(1,34.1) = 0.30, p = 0.59; Backward: F(1, 34) = 2.29, p = 0.14) (*Figure 4B*). There were no interactions between perturbation magnitude and group (Forward: F(2, 6.38) = 1.59, p = 0.21; Backward: F(2, 68) = 0.39, p = 0.68).

Individuals with PD exhibited a larger acceleration driven component of destabilizing muscle activity compared to OAs. We further decomposed the output from the destabilizing feedback loop to antagonist muscles (*Figure 4A* red trace) into its component parts attributed to the CoM acceleration feedback component (k_aD_) and the CoM velocity and displacement feedback component (k_vD_+k_dD_). Individuals with PD had larger k_aD_ component compared to OAs in forward perturbations (Forward: t(34) = -2.05, p = 0.048; Backward: t(34) = 1.174, p = 0.25). There was no difference between OA and PD groups in the contribution of the k_vD_+k_dD_ component in either perturbation direction (Forward: (t(34) = 0.86, p = 0.40; Backward (t(34) = 1.37, p = 0.18).

Contributions from the stabilizing feedback loop to antagonist muscles (*Figure 4A* green trace) that occurred at the end of the ramp-and hold perturbations increased with perturbation magnitude, but only the backward perturbation direction (Forward: F(1, 63.9) = 0.15, p = 0.86; Backward: F(2, 68) = 3.98, p = 0.023) (*Figure 4B*). There was no effect of group on the output from the stabilizing feedback loop (Forward: F(1, 33.9) = 0.95, p = 0.34; Backward: F(1, 34) = 2.20, p = 0.15) (*Figure 4B*). There were no interactions between perturbation magnitude and group (Forward: F(2, 63.9) = 1.41, p = 0.25; Backward: F(2, 68) = 0.21, p = 0.81).

### Destabilizing antagonist co-contraction occurred at latencies consistent with higher-order mediation

The onset of the antagonist destabilizing feedback loop occurred at an intermediate latency between the onset of the agonist LLR1 and LLR2. The antagonist destabilizing co-contraction had an onset latency (λ_D_) of 180 ± 3.6ms in OAs and 173 ± 2.9ms for individuals with PD. There was no difference in onset latency between OA and PD groups (Forward: t(34) = 1.76, p = 0.09; Backward: t(34) = -0.80, p = 0.43). The onset of the antagonist destabilizing co-contraction occurred at 119 ± 1.6ms in OAs and 116 ± 1.3ms in people with PD, after the initial burst of agonist muscle activity (λ_1_), but prior to the second burst of agonist muscle activity (λ_2_) at 200 ± 3.2ms in OAs and 215 ± 3.9ms in people with PD.

### Correlations to clinical measures

The output from the destabilizing feedback loop was negatively correlated with miniBEST scores in the OA group for both perturbation directions (Forward: R^2^ = 0.69, p = 0.043; Backward: R^2^ = 0.70, p = 0.045). Individuals with PD showed an opposite trend, as there was a positive relationship between the output from the antagonist destabilizing feedback loop and higher miniBEST scores; this trend was only present in the forward perturbations and did not reach significance (Forward: R^2^ = 0.48, p = 0.083; Backward: R^2^ = 0.40, p = 0.90).

There was a negative trend between the contribution from the longer-latency feedback loop and miniBEST in OA, but not PD. However, this trend did not reach significance in OAs (Forward: R^2^ = 0.63, p = 0.088; Backward: R^2^ = 0.33, p = 0.25) or individuals with PD (Forward: R^2^ = 0.53, p = 0.55; Backward R^2^ = 0.24, p = 0.91).

None of the assessments of PD motor symptom severity (MDS UPDRS-III score, PD phenotype, PIGD-subscore, Hoehn and Yahr state) nor PD duration could explain the variance in any of the components identified by our neuromechanical models (*Table 2*).

**Table 2:** Statistical comparisons of neuromechanical model outputs and Parkinson’s disease specific clinical measures.

| Integrated Component | MDS-UPDRS III | PD Phenotype | PIGD-subscore | Hoehn & Yahr state | PD Duration |
| --- | --- | --- | --- | --- | --- |
| Agonist short-latency | Forward:<br>$p = 0.85$<br>Backward:<br>$p = 0.84$ | Forward:<br>$F(2,14.1)=0.4$,<br>$p=0.68$<br>Backward:<br>$F(2,14)=1.16$, $p = 0.34$ | Forward:<br>$F(6, 9.53) = 2.02$, $p = 0.16$<br>Backward:<br>$F(6,10) = 0.85$, $p = 0.56$ | Forward:<br>$F(3,12.3)=3.5$ ,<br>**$p=0.049^*$**<br>Backward:<br>$F(3,13)=0.17$ , p=0.92 | Forward:<br>$p = 0.87$<br>Backward:<br>$p = 0.89$ |
| Agonist long-latency | Forward:<br>p = 0.46<br>Backward:<br>p = 0.97 | Forward:<br>F(2,14.2)=0.4,<br>p=0.68<br>Backward:<br>F(2,14)=1.0,<br>p=0.38 | Forward:<br>F(6,10.1) = 1.26,<br>p = 0.36<br>Backward:<br>F(6,10) = 0.51, p<br>= 0.79 | Forward:<br>F(3,12.8)=1.0,<br>p=0.41<br>Backward:<br>F(3,13)=0.5,<br>p=0.69 | Forward:<br>p = 0.83<br>Backward:<br>p = 0.83 |
| Antagonist stabilizing | Forward:<br>p = 0.19<br>Backward:<br>p = 0.75 | Forward:<br>F(2,14.2)=1.21,<br>p=0.33<br>Backward:<br>F(2,14)=0.29,<br>p=0.75 | Forward:<br>F(6, 9.41) =<br>0.84, p = 0.57<br>Backward:<br>F(6,10) = 0.55, p<br>= 0.76 | Forward:<br>F(3,12.8) = 0.37, p<br>= 0.77<br>Backward:<br>F(3,13) = 0.17, p =<br>0.91 | Forward:<br>p = 0.59<br>Backward:<br>p = 0.29 |
| Antagonist destabilizing | Forward:<br>p = 0.59<br>Backward:<br>p = 0.44 | Forward:<br>F(2,13.9)=0.3,<br>p=0.74<br>Backward:<br>F(2,14)=0.15,<br>p=0.86 | Forward:<br>F(6,9.92) = 1.31,<br>p = 0.34<br>Backward:<br>F(6,10) = 0.1, p<br>= 0.995 | Forward:<br>F(3,12.8)=1.2,<br>p=0.35<br>Backward:<br>F(3,13)=0.10,<br>p=0.96 | Forward:<br>p = 0.75<br>Backward:<br>p = 0.23 |
| Antagonist destabilizing (kaD component) | Forward:<br>p = 0.90<br>Backward:<br>p = 0.39 | Forward:<br>F(2,13.8)=0.55,<br>p=0.59<br>Backward:<br>F(2,14)=1.05,<br>p=0.38 | Forward:<br>F(6, 9.42) =<br>0.74, p = 0.63<br>Backward:<br>F(6,10) = 0.66, p<br>= 0.69 | Forward:<br>F(3,12.6) = 0.69, p<br>= 0.57<br>Backward:<br>F(3,13) = 0.28, p =<br>0.84 | Forward:<br>p = 0.47<br>Backward:<br>p = 0.37 |
| Antagonist destabilizing (kvD+kdD component) | Forward:<br>p = 0.59<br>Backward:<br>p = 0.50 | Forward:<br>F(2,14)=0.34,<br>p=0.72<br>Backward:<br>F(2,14)=0.27,<br>p=0.77 | Forward:<br>F(6, 9.97) =<br>1.31, p = 0.34<br>Backward:<br>F(6,10) = 0.07, p<br>= 0.998 | Forward:<br>F(3,12.9) = 1.14, p<br>= 0.37<br>Backward:<br>F(3,13) = 0.1, p =<br>0.96 | Forward:<br>p = 0.75<br>Backward:<br>p = 0.24 |

## DISCUSSION

Overall, this work demonstrates that a neuromechanical model can decompose perturbation evoked muscle activity into hierarchical components attributed to different neural substrates based on latency that are relevant to clinical measures of balance ability. Our data support the hypothesis that parallel sensorimotor feedback loops engaging hierarchical circuits contribute to perturbation evoked muscle activity during reactive balance recovery in OAs with and without PD. These results provide experimental and computational evidence that longer-latency agonist muscle activity associated with the LLR2 can be explained by sensory information encoding CoM kinematics at delays consistent with transcortical feedback loops in OAs with and without PD. We further show that cortically-mediated responses increase with balance challenge in OAs with and without PD. Furthermore, in an exploratory comparison we show that the LLR2 is recruited at smaller perturbation magnitudes in OAs than YAs (36), regardless of PD status (*Figure 5*). Our findings are in consistent with the compensation-related utilization of neural circuits hypothesis (CRUNCH) (58,59), as OAs have increased activation of cortical regions compared to YAs, to compensate for declining efficiency of processing in subcortical circuits. We also show that perturbation-evoked destabilizing antagonist activity occurs at latencies consistent with higher-order mediation. Both OAs and people with PD exhibited similar amounts of destabilizing antagonist muscle activity that occurred at similar onset latencies. The magnitude of this destabilizing component was correlated with an individual’s balance ability in OAs, but not individuals with PD. This implies that neuromechanical modeling of reactive balance control may provide a useful tool to index individual differences in the hierarchical control of balance, which may help clinicians optimize future rehabilitation training (60,61).

**Figure 5:**
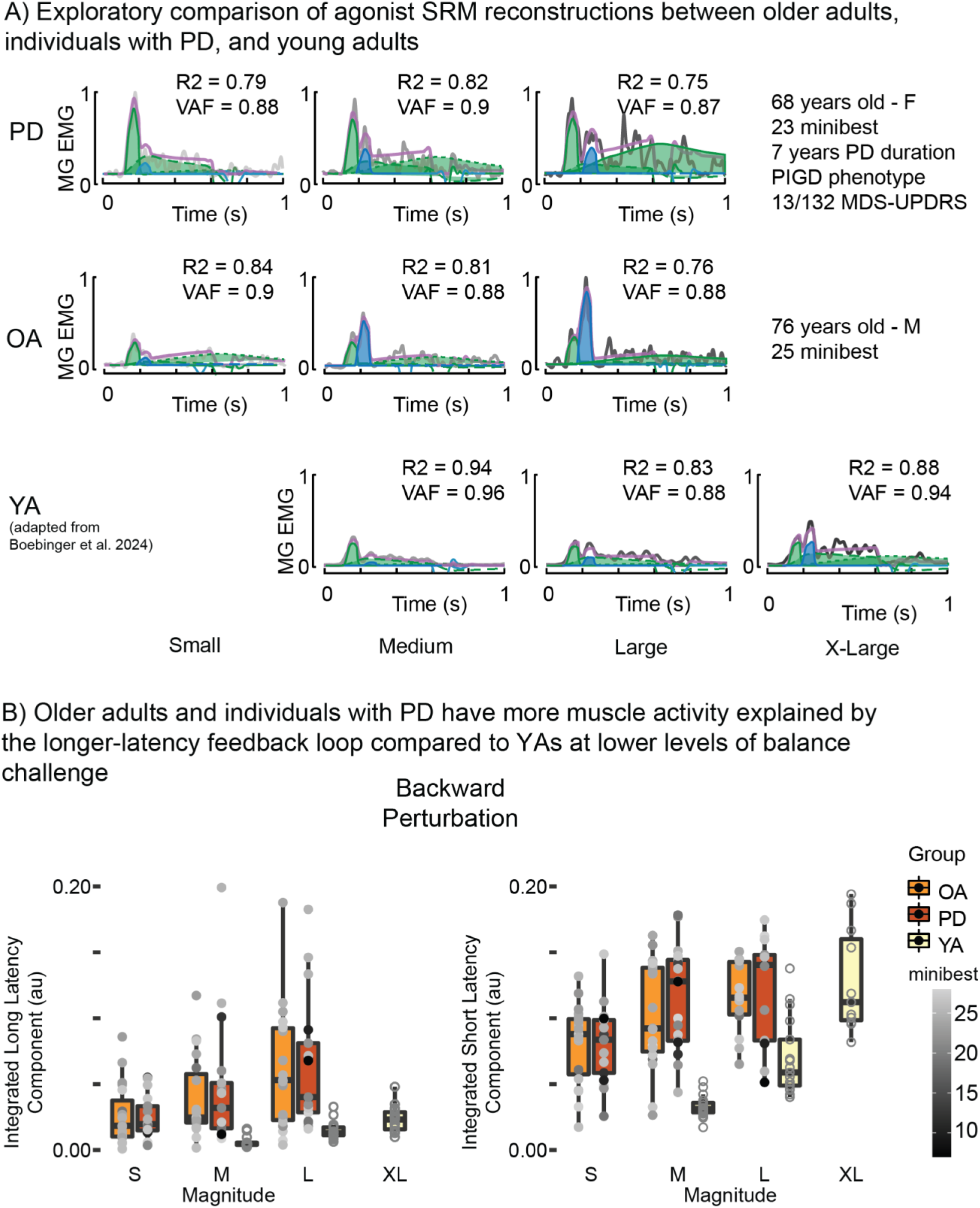
Exploratory comparison to previously published data in young adults (YAs). We compared the outputs from the shorter- and longer-latency feedback loops of the agonist neuromechanical model (Figure 3A). Comparisons between groups are qualitative due to differences in the experimental paradigms between the YAs and the OAs with and without PD.

Our neuromechanical model could provide a more mechanistic assessment of cortical engagement during balance control. Our current understanding of cortical contributions to balance in aging and PD is typically either inferred through dual task interference analysis (1,10,18–20) or measures of cortical oxygen metabolism (13,14,21,22). For dual task interference assessments, cortical engagement is typically inferred when primary motor task, i.e. standing balance, is performed concurrently with a secondary cognitive task, i.e. mental math (10,18,62). Decreased balance performance when the secondary task is performed concurrently to primary task are thought to arise because the secondary task diverts cortical resources, defined as the brain’s task-general information processing capacity that is shareable across concurrent tasks in a graded manner (63–65), away from primary task thereby impacting performance. Cortical engagement has also been assessed through optical measurements of cortical oxygen metabolism using functional near-infrared spectroscopy (fNIRS) (13,14,21,22). However, fNIRS is often only used to assess prefrontal oxygen metabolism as hair on the rest of the scalp prevents this technique from being used on areas over other neural structures (66–70). Both techniques lack temporal resolution due to inherent delay between neurovascular coupling in the case of fNIRS or between muscle activity and movement in the case of behavioral assessments. Therefore, how these findings will be reflected in muscle activity was unable to be determined. Assessments of perturbation evoked muscle activity like those uses here potentially offer a more robust way to characterize cortical contributions to balance control as muscle activity has millisecond-level temporal resolution which is necessary due to the discrete nature of balance correcting responses.

Our results are consistent with prior work that demonstrates increased cortical engagement during balance with aging and impairment (1,10,11,13,14,18–20). Older adults exhibit increased activation of cortical areas relative to younger adults performing similar tasks (58,59). Furthermore, individuals with impaired balance such as in PD exhibit further increases in cortical activity relative to unimpaired OAs (1,10,13,14,18). In an exploratory analysis, we compared the reconstructions of agonist muscle activity from OAs with and without PD to previously published reconstructions from YAs (36) (*Figure 5*). OAs, regardless of PD, exhibit a component of agonist muscle activity that could be explained by CoM kinematics at a delay consistent with transcortical feedback loops, even at the smallest perturbation magnitude (5.1 cm displacement), whereas for young adults this muscle activity is largely absent at a larger magnitude (7.7cm displacement) (*Figure 5*). There are some notable differences between the perturbation paradigm used here and those used previously in the YA group which prevent direct comparisons. Namely, the YA paradigm only had perturbations in a single direction while the OA and PD paradigm had perturbations in both directions. Therefore, the YA group would have been more able to adapt their motor response to the perturbation paradigm, which has been shown to alter our neuromechanical model’s parameters (34), since adaptation could lead to balance correcting responses to be more automatic and thereby require less cortical input.

Contrary to our hypothesis that individuals with PD will have increased involvement of higher-order circuits during balance control, we found that OAs with PD had similar amounts of longer-latency muscle activity as OAs without PD. Perturbation evoked responses progressively engage higher order centers at increasing latency (11,27). The initial burst of agonist muscle activity (LLR1) occurred at latencies around 120ms, consistent with prior iterations of this neuromechanical model (15,28,29,34,36). Muscle activity at this latency are considered to be mediated by brainstem sensorimotor circuits (11) as evidenced by decerebrate animals being able to maintain balance and exhibit intact, perturbation-specific patterns of muscle activation when exposed to balance perturbations (9). Additionally, in humans, perturbation evoked cortical activity occurs simultaneously with the LLR1, and therefore cannot contribute to this aspect of muscle activity (36,55,71). A second burst of agonist muscle activity (LLR2) occurred at a latencies around 200ms, consistent with the latencies found in prior iterations of this model (36). Muscle activity at this longer latency has the potential to be mediated by higher-order, presumably transcortical circuits (11,25), as this activity can be altered by transcranial magnetic stimulation, while shorter latency muscle activity cannot (25,72). Additionally, the latency of the LLR2 was about twice that of perturbation-evoked cortical activity recorded via electroencephalography in YAs (36), making it possible for evoked cortical activity to contribute to this aspect of balance correcting muscle activity.

Our findings in reconstructions of antagonist muscle activity corroborate previous findings showing that antagonist co-contraction can be explained by sensorimotor transformations of CoM kinematic errors at delays consistent with input from higher-order neural circuits and is related to balance ability (15). The onset of the destabilizing muscle activity occurred around 180ms in both the OA and PD groups, similar to previous implementations of this model (15). Interestingly, this destabilizing co-contraction occurs after the shorter-latency, presumably brainstem-mediated, agonist muscle activity. The initial burst of agonist muscle activity is thought to be governed by reticulospinal circuits (33) and occurred ∼120ms for both groups. The onset of the destabilizing antagonist muscle activity was substantially longer (∼60ms on average), which is sufficiently long for basal ganglia involvement (15). Additionally, the onset of destabilizing antagonist co-contraction occurred prior to the onset of the longer-latency agonist muscle activity identified by our neuromechanical model. This could suggest that these two components of perturbation evoked muscle activity are mediated by different neural circuits. However, we can only speculate on the neuroanatomical substrates involved in generating motor output since we lack neuroimaging techniques and rely on the latency of muscle activity. It is important to note that while we find similar results, we only found this phenomenon in the forward perturbations, whereas the previous publication was focused on backward perturbations. This discrepancy could result from differences in the experimental paradigm between these studies. Here examined 3 perturbation magnitudes while previously only a single magnitude was used. Furthermore, we are examining data from seventeen individuals with PD whereas the previous study examined data collected from forty-four individuals with PD. The differences in experimental paradigm and number of participants may explain discrepancies between what we show here and what was previously published.

Recordings of muscle activity during reactive balance recovery coupled with our neuromechanical model could offer a means to index shifts in the hierarchical control of balance, which would be useful for tracking changes in balance control throughout rehabilitation interventions and assessing the efficacy of balance-assisting devices. Reactive balance paradigms could be used to quantify the LLR1 and LLR2 components and thereby provide a metric of cortical contributions to balance control without the need for measures of brain activity. Changes in these hierarchical components could then be tracked throughout disease progression and rehabilitation interventions at different levels of balance challenge. Additionally, the effect of assistive devices on an individual’s control of balance could be inferred by the degree to which the balance control while using a device is cortically-mediated. Furthermore, our neuromechanical modeling technique can be extended to reconstruct joint torques, which would circumvent the need to record muscle activity (73).

## DATA AVAILABILITY

All data, model code, and statistical analysis code can be found online at: https://osf.io/56u43/?view_only=d82dd6b36d5f4313ada65f4ec6203d6c

## DISCLAIMERS

The authors have no disclaimers

## AUTHOR CONTRIBUTIONS

L.H.T and A.M.P Conceived and designed research, A.M.P performed experiments, S.E.B and J.X analyzed data, S.E.B, G.M, A.M.P, J.L.M, M.R.B, and L.H.T interpreted results of experiments, S.E.B prepared figures, S.E.B drafted manuscript, S.E.B, J.X, G.M, A.M.P, J.L.M, M.R.B, and L.H.T edited and revised manuscript, S.E.B, J.X, G.M, A.M.P, J.L.M, M.R.B, and L.H.T approved final version of manuscript.

